# Transplacental innate immune training via maternal microbial exposure: the XBP1-ERN1 axis in programming dendritic cell precursors

**DOI:** 10.1101/848242

**Authors:** Kyle T. Mincham, Anya C. Jones, Marie Bodinier, Naomi M. Scott, Jean-Francois Lauzon-Joset, Philip A. Stumbles, Anthony Bosco, Patrick G. Holt, Deborah H. Strickland

## Abstract

We recently reported that the offspring of mice treated during pregnancy with the microbial-derived immunomodulator OM-85 manifest striking resistance postnatally to allergic airways inflammation, and localised the potential treatment target to the fetal cDC progenitor compartment which expands to increase the pool of precursors available at birth, enabling accelerated postnatal seeding of the lung mucosal cDC network required for establishment of immunological homeostasis in the airways. Here, we profile maternal OM-85 treatment-associated transcriptomic signatures in fetal bone marrow, and identify a series of immunometabolic pathways which provide essential metabolites for accelerated myelopoiesis, that are hallmarks of classical “immune training”. In addition, the cDC progenitor compartment displayed treatment-associated activation of the XBP1-ERN1 signalling axis which has previously been shown to be essential for tissue survival of cDC, particularly within the lung microenvironment. Our forerunner studies indicate uniquely rapid turnover of airway mucosal cDCs at baseline, with further large-scale upregulation of population dynamics during aeroallergen and/or pathogen challenge. XBP1-ERN1 signalling plays a key role in mitigation of ER stress-associated toxicity which frequently accompanies DC hyper-activation during intense immunoinflammatory responses, and we suggest that enhanced capacity for XBP1-ERN1-dependent cDC survival within the airway mucosal tissue microenvironment may be a crucial element of the OM-85-mediated transplacental “innate immune training” process which results in enhanced resistance to airway inflammatory disease during the high-risk early postnatal period.

## Introduction

The neonatal period represents a time of high risk for infection-related morbidity/mortality resulting from the combined effects of maturational deficiencies in both anti-microbial defense mechanisms that mediate pathogen recognition and elimination, and in the accompanying regulatory mechanisms required for calibration of these responses to minimise inflammatory damage to host tissues^1, 2^. With respect specifically to the lung tissue microenvironment, a crucial factor governing the kinetics of postnatal acquisition of immune competence is the rate of development of the airway mucosal cDC network which controls local immune surveillance^3, 4^.

In addition to increased susceptibility to infectious diseases, the seeds for development of a range of non-communicable diseases exemplified by asthma and aero-allergies are also frequently sewn during this early postnatal window period^5^, suggesting that maturational deficiencies in immune function(s) may also be risk factors in this context. In this regard, maternal immune perturbations have been acknowledged to significantly influence fetal immune development, but the underlying mechanisms remain poorly characterised^6^. Emerging epidemiological data from studies on traditional farming families in Europe and USA suggesting that benign environmental microbial exposures of mothers during pregnancy can promote prenatal immune maturation within their offspring, leading to reduced susceptibility to postnatal development of respiratory inflammatory diseases^7, 8^, have stimulated wide-spread interest in this issue. This capacity for microbial exposures to modulate immune development is consistent with the paradigm of “immune training”, whereby exposure to certain classes of microbial stimuli can alter the long-term functional state of innate immune cells, occurring at the progenitor level in the bone marrow (BM)^9-11^, leading to optimised peripheral immune responsiveness to other unrelated microorganisms^12^. With this in mind, there is growing interest in the concept that immune training can be therapeutically harnessed^13^, particularly during prenatal development, to enhance immunocompetence within the offspring^14^.

We recently reported that oral treatment of pregnant mice with the microbial-derived immunomodulatory agent OM-85 reduces susceptibility of their offspring to the development of Th2-driven allergic airways inflammation, and identified myeloid progenitors in the offspring BM (which supply precursor DC to eventually populate mucosal DC networks) as a major target for maternal treatment effects^15^. In the study presented here, we employed transcriptomic profiling to characterise gene networks activated in fetal BM (fBM) as a result of maternal OM-85 treatment, and identify the principal treatment targets as immunometabolic pathways supplying cellular cholesterol essential for rapid expansion of myeloid precursor compartments, and which have previously been recognised as hallmarks of classical immune training-associated gene signatures. We additionally identify a specific pathway in the cDC precursor compartment which has previously been associated with survival-under-stress, especially within the lung mucosal microenvironment.

## Results

To elucidate the mechanisms-of-action of maternal OM-85 treatment, we examined the *in utero* fetal response at gestation day 18.5 (Figure 1A), 2 days prior to expected natural term delivery.

**Figure 1.**
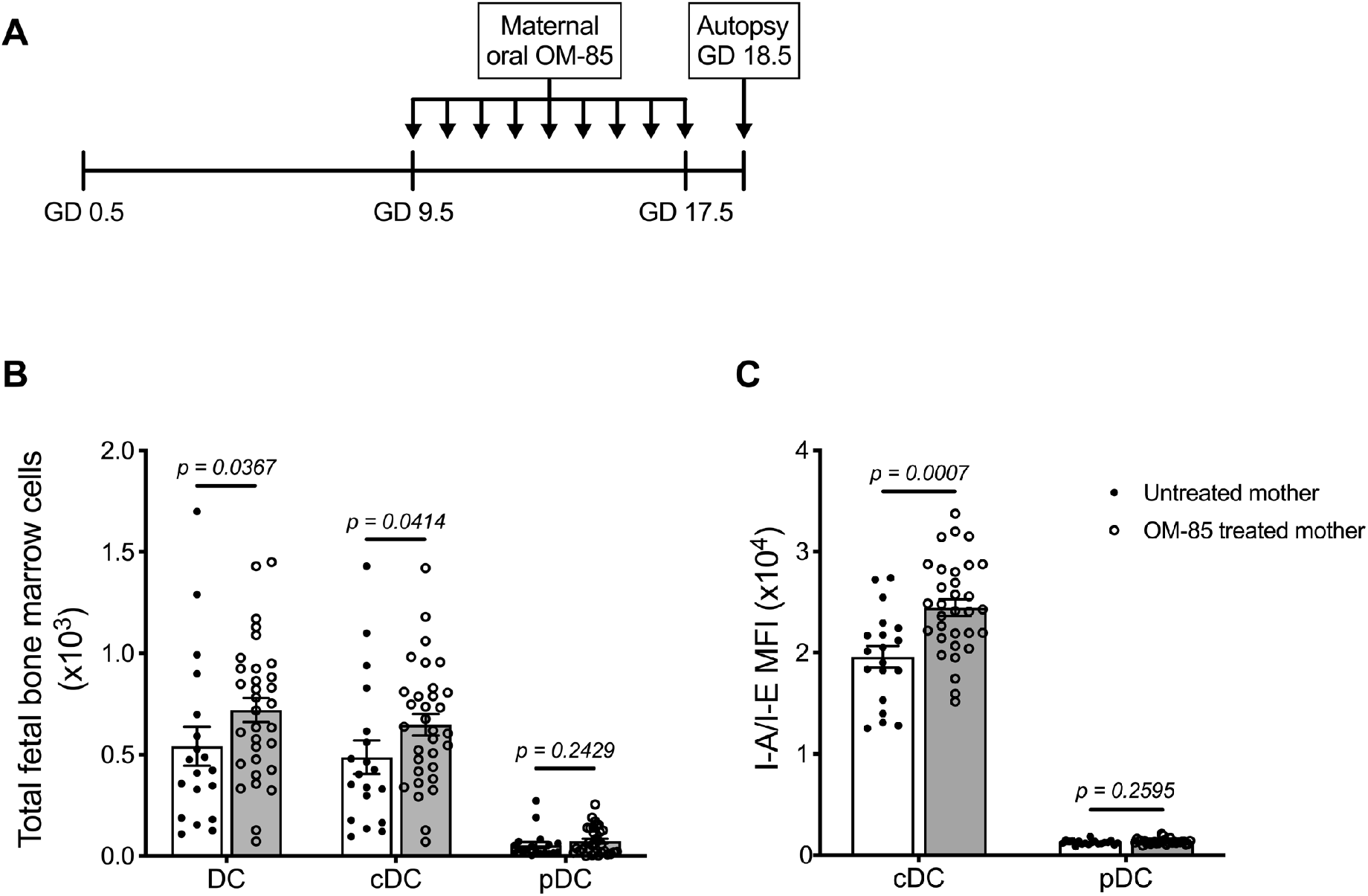
Maternal OM-85 treatment during pregnancy selectively modulates the fetal bone marrow conventional DC subset. **(A)** Kinetics of maternal OM-85 treatment beginning at gestation day (GD) 9.5 with daily oral treatment until GD 17.5 and autopsy 24 hours post final treatment at GD 18.5. **(B)** Absolute numbers of dendritic cells (DC), CD11b^+^B220^-^CD11c^+^Gr-1^-^ SIRP-α^+^I-A/I-E^+^ conventional dendritic cells (cDC) and CD11b^-^B220^+^CD11c^+^Gr-1^+^I-A/I-E^+^ plasmacytoid dendritic cells (pDC) in BM of fetuses from OM-85-treated and untreated mothers. **(C)** Mean fluorescence intensity (MFI) of I-A/I-E expression on cDCs and pDCs in fBM. Data are presented from individual animals comparing fetuses from OM-85-treated and untreated mothers and displayed as bar graphs showing mean ± SEM of *n* = 8 independent experiments. Statistical significance was determined using Mann-Whitney *U* test (B) or Student’s *t* test (C) based on distribution of the data as determined by D’Agostino-Pearson omnibus normality test.

### Maternal OM-85 treatment selectively accelerates functional maturation of cDCs in fetal bone marrow

Phenotypic analysis of the fBM myeloid compartment using multicolour flow cytometry revealed significant expansion of the total dendritic cell (DC) pool in fetuses from OM-85 treated mothers as compared to equivalent fetal samples from untreated mothers (Figure 1B). Further characterisation of the BM DC response demonstrated that this increase was restricted to the conventional DC (cDC) subset as previously described^15^, with no parallel changes observed in plasmacytoid DC (pDC) (Figure 1B). These cDC-specific changes in fBM mirror our recent observations of increased cDC yields from BM cultures and peripheral lung from the offspring of OM-85-treated mothers in the early postnatal period^15^. We next turned our attention to fetal DC maturation state as determined by surface I-A/I-E expression. As shown in Figure 1C, maternal OM-85 treatment for the last half of gestation enhanced I-A/I-E expression on cDC in fBM when compared to fBM cDC from untreated mothers. Collectively, these observations suggest that transplacental signals generated at the feto-maternal interface following OM-85 treatment during pregnancy can “train” the developing fetal immune system via accelerating the development of functional competence within the fBM cDC compartment.

### Expansion of fetal bone marrow myeloid progenitor subsets following maternal OM-85 treatment

Previous studies from our laboratory have additionally identified postnatal expansion of BM myeloid progenitor (MP) cell populations as an effect of maternal treatment with OM-85 during pregnancy^15^. These findings mirror that of recent studies which identified modulation of BM MP as an important component of conventional immune training mediated by both β-glucan-^10^ and Bacillus Calmette-Guérin (BCG)^11^. Based on these findings, we hypothesised that the enhanced cDC population within fBM following maternal OM-85 treatment would also be accompanied by concomitant upregulation of upstream MP subsets. Flow cytometric analysis of fBM (Figure 2A) Lin^-^IL-7Rα^-^c-Kit^+^Sca-1^-^ MP (Figure 2B), Lin^-^IL-7Rα^-^c-Kit^+^Sca-1^-^ CD16/32^hi^CD34^+^ granulocyte-macrophage progenitor (GMP; Figure 2C) and Lin^-^IL-7Rα^-^c-Kit^+^Sca-1^-^CD16/32^hi^CD34^+^CX3CR1^+^Flt-3^+^ macrophage-dendritic cell progenitor (MDP; Figure 2D) populations demonstrated a total increase in all 3 progenitor subsets within the BM compartment following maternal OM-85 treatment, when compared to fBM from untreated mothers. Consistent with the findings in Figure 1B and C, these data provide further evidence that maternal OM-85 treatment selectively modulates the offspring BM myeloid lineage *in utero*, beginning at the early-stage myeloid progenitor level through to the terminal cDC populations which are responsible for seeding peripheral tissues during early postnatal life to provide local DC-mediated immune surveillance.

**Figure 2.**
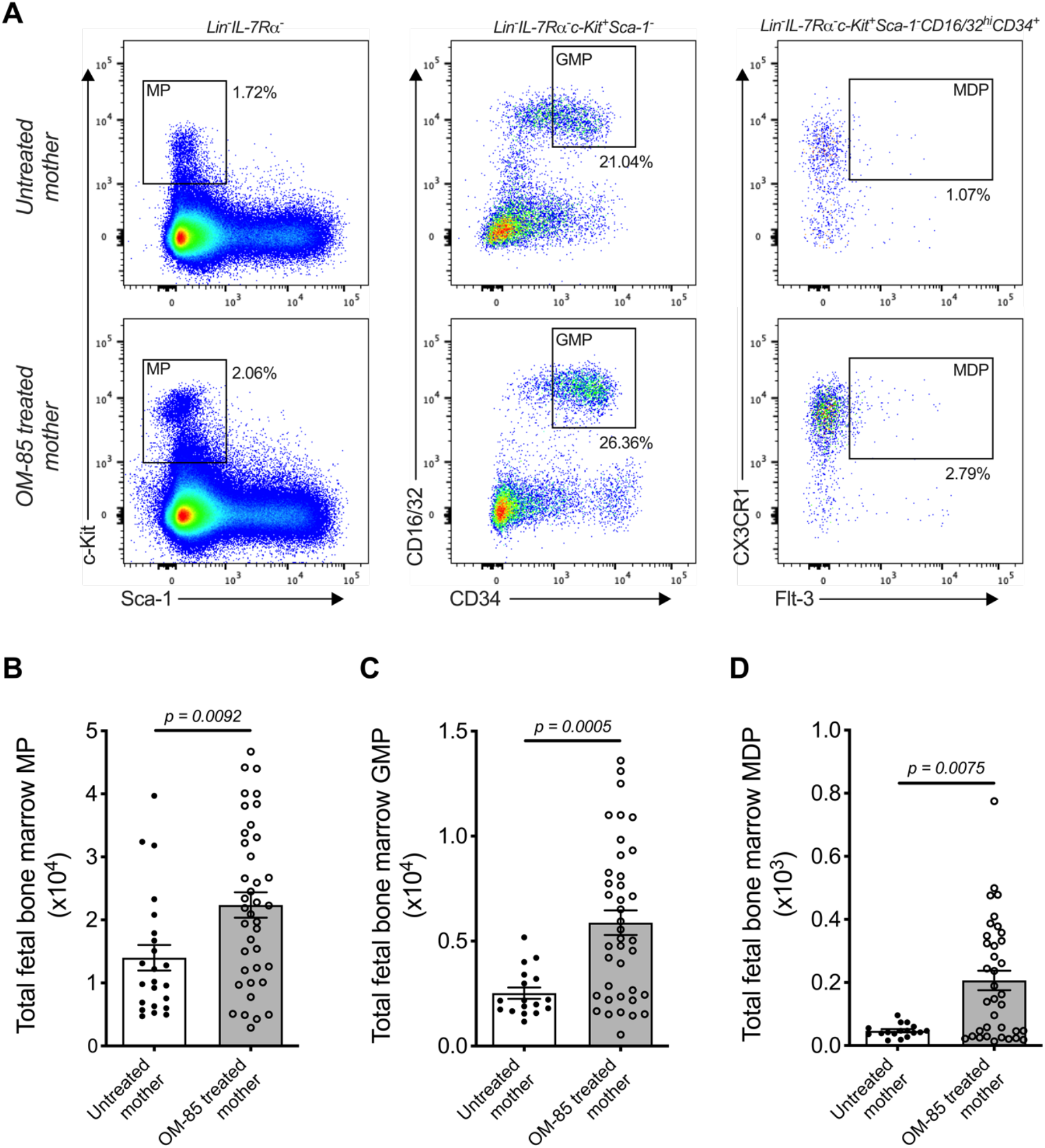
Treatment of mothers with OM-85 during pregnancy boosts myeloid progenitor subsets within fetal bone marrow. **(A)** Representative flow cytometry plots for the identification of Lin^-^IL-7Rα^-^c-Kit^+^Sca-1^-^ myeloid progenitors (MP), Lin^-^IL-7Rα^-^c-Kit^+^Sca-1^-^ CD16/32^hi^CD34^+^ granulocyte-macrophage progenitors (GMP) and Lin^-^IL-7Rα^-^c-Kit^+^Sca-1^-^ CD16/32^hi^CD34^+^CX3CR1^+^Flt-3^+^ macrophage-dendritic cell progenitors (MDP) within fBM. Absolute numbers of **(B)** MP, **(C)** GMP and **(D)** MDP in BM of fetuses from OM-85-treated and untreated mothers. Data are presented from individual animals comparing fetuses from OM-85-treated and untreated mothers and displayed as bar graphs showing mean ± SEM of *n* = 8 independent experiments. Statistical significance was determined using Student’s *t* test (C) or Mann-Whitney *U* test (B, D) based on distribution of the data as determined by D’Agostino-Pearson omnibus normality test.

### Maternal OM-85 treatment activates key regulators of the UPR pathway in fetal bone marrow

To gain further insight into the molecular mechanisms underpinning the maternal OM-85-treatment effects, we employed transcriptomic profiling of fBM cells. Comparison of the transcriptomic profiles in the treated versus untreated groups indicated that maternal OM-85 treatment resulted in 152 differentially expressed genes (DEG) in fBM (119 upregulated, 33 downregulated; Figure 3A, Supplementary Table 1). We then interrogated the DEG for enrichment of biological pathways employing *InnateDB*^16^, focusing on the upregulated DEG response given the limited number of downregulated DEG identified. In fBM, upregulated DEG were enriched for genes involved in multiple aspects of protein metabolism, the endoplasmic reticulum (ER) stress response, the unfolded protein response (UPR), cholesterol biosynthesis and lipid metabolism (Supplementary Table 2).

**Figure 3.**
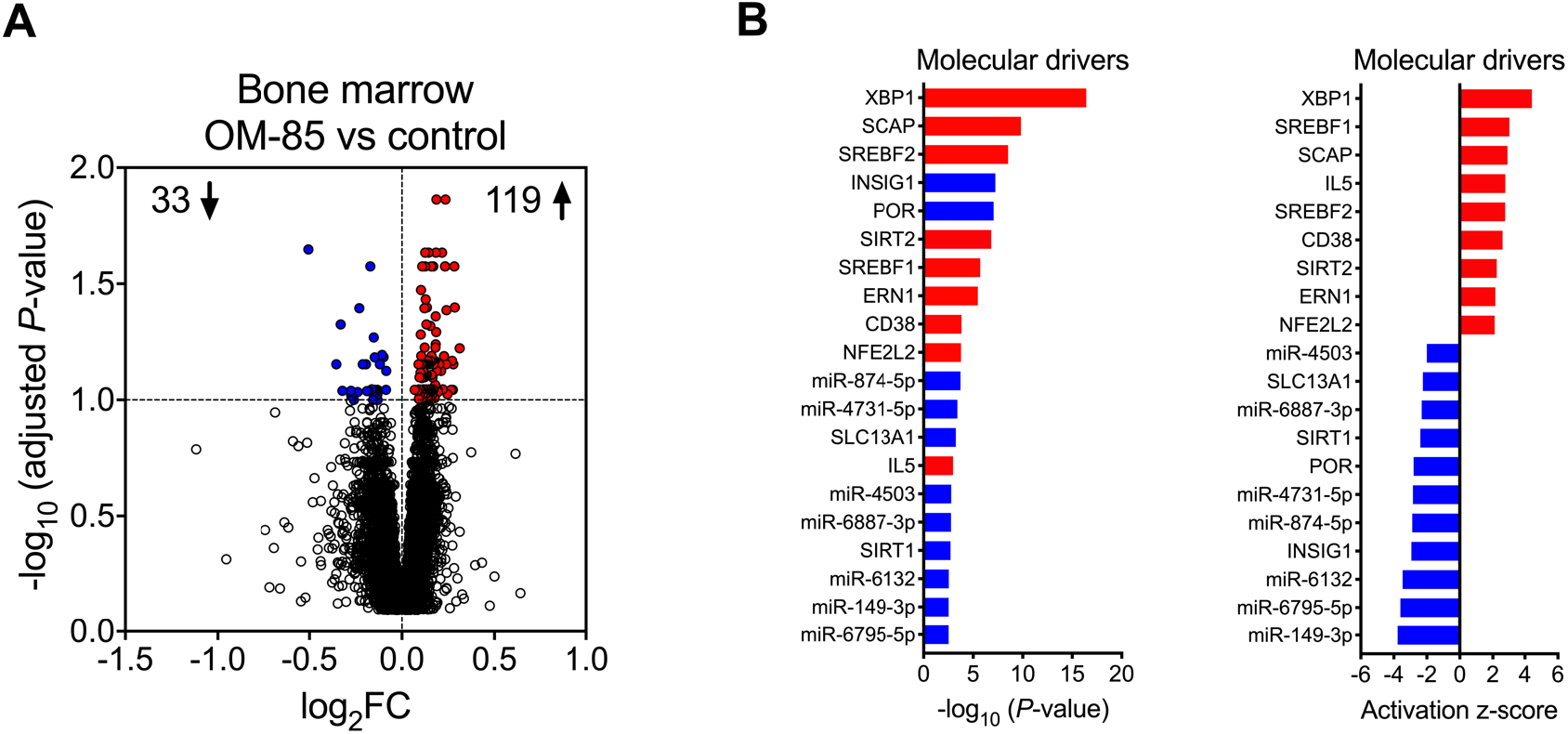
Maternal OM-85-induced changes in fetal bone marrow gene expression profiles. **(A)** Differentially expressed genes (DEG) within fBM comparing offspring from OM-85-treated and untreated mothers. DEG are summarised as a volcano plot showing data along axes of statistical significance (-log_10_ adjusted *P*-value) and differential expression magnitude (log2 fold change) for *n* = 16 individual animals per group collected from *n* = 7 independent experiments. Dashed horizontal lines indicate a False Discovery Rate (FDR) adjusted *P*-value < 0.1. Genes shown in red were upregulated and those shown in blue were downregulated. **(B)** activated (red) and inhibited (blue) molecular drivers of the differential expression patterns were identified using Upstream Regulator Analysis.

Upstream regulator analysis was then performed to identify putative molecular drivers of all observed DEG. The data revealed X-box binding protein 1 (XBP1), a transcription factor central to the UPR^17^ and crucial in the development, survival and function of multiple cell types including plasma cells^18^, eosinophils^19^, natural killer (NK) cells^20^, T-cell subsets^21^ and DCs^22, 23^, as the most strongly activated molecular driver within fBM associated with maternal OM-85 treatment effects (*P*-value = 3.81×10^-17^, Z-score = 4.427, Figure 3B; Supplementary Table 3), and consistent with this, its downstream target, the canonical UPR sensor Activating Transcription Factor 6 beta (ATF6β)^24, 25^ was upregulated (Supplementary Table 1 and 3). Additionally, Endoplasmic Reticulum To Nucleus Signalling 1 (ERN1) was identified as an activated driver gene within the fBM following maternal OM-85 treatment (*P*-value = 3.32×10^-6^, Z-score = 2.156; Figure 3B; Supplementary Table 3). Identification of ERN1 is crucial given that during the ER stress response, this gene encodes the ER stress sensor protein inositol-requiring enzyme 1 (IRE1α), responsible for the unconventional cleavage of a 26 nucleotide fragment from *Xbp1* mRNA, resulting in the generation of the active spliced variant of XBP1 (XBP1s) and enabling it to function as a potent transcription factor within the UPR signalling pathway^25^. Collectively, these findings suggest that activation of the UPR pathway may be a central component of the immune training mechanism induced in fBM as a result of maternal OM-85 treatment. In addition to UPR pathway drivers, maternal OM-85 treatment also resulted in the upstream activation of multiple drivers central to immunometabolic pathways involved in cellular cholesterol homeostasis, including Sterol Regulatory Element Binding Transcription Factor 1 (SREBF1; *P*-value = 1.95×10^-6^, Z-score = 3.056) and 2 (SREBF2; *P*-value = 2.95×10^-9^, Z-score = 2.745) and Sterol Regulatory Element Binding Protein Cleavage-Activating Protein (SCAP; *P*-value = 1.52×10^-10^, Z-score = 2.949; Figure 3B; Supplementary Table 3). This was associated with upregulation of their downstream target Low-Density Lipoprotein Receptor (LDLR; Supplementary Table 1 and 3), while Insulin Inducible Gene 1 (INSIG1) was inhibited in fBM following maternal OM-85 treatment (*P*-value = 5.81×10^-8^, Z-score = −2.931; Figure 3B; Supplementary Table 3).

Additional candidate drivers in fBM following maternal OM-85 treatment included CD38, IL5, and an array of microRNAs (miR) (Figure 3B; Supplementary Table 3) recognised principally in the context of cancer-associated functions^26-28^. Of note, miR-149-3p (strongly downregulated in Figure 3B) has been shown to negatively regulate Toll-like receptor (TLR) 4 expression in murine monocytic cells *in vitro^29^* and it is possible that other miRs may have similar (but as yet undefined) innate immune regulatory functions^30^. The relevance of finding TLR4 upregulation in BM-derived myeloid cells in this model merits further investigation. Likewise, the identification of the T-cell activation-associated markers CD38 and IL-5 suggests possible contributions from activated T-cells, and these possibilities will also be addressed in follow up studies.

### Upregulation of XBP1s expression is restricted to fetal bone marrow cDC precursors following maternal OM-85 treatment

Finally, to obtain evidence confirming that the activated form of XBP1 was upregulated, we measured expression of the active spliced variant of XBP1 (XBP1s) at the protein level within fBM. Using multicolour flow cytometry, we identified significant upregulation of fBM CD11b^+^CD11c^+^ pre-cDCs expressing intracellular XBP1s following maternal OM-85 treatment (Figure 4A, B). While intracellular XBP1s expression was additionally localised within fetal CD19^+^B220^+^ B cells, NKp46^+^CD11b^+^B220^+^CD11c^lo^ NK cells and CD3^+^ T-cells (Supplementary Figure 1), as previously described in the literature^18, 20, 21, 31^, maternal OM-85 treatment had no impact on XBP1s expression levels in these cell types.

**Figure 4.**
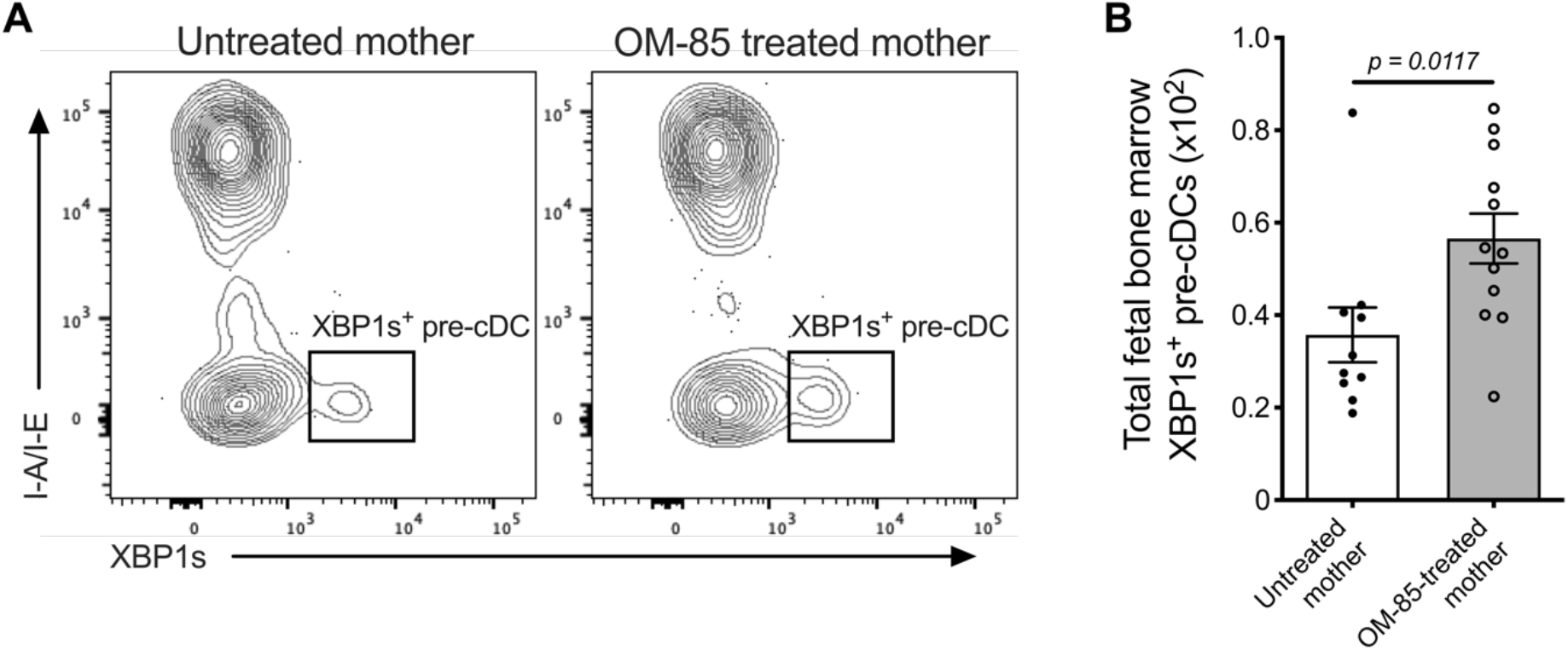
XBP1s expression in pre-cDCs within fetal bone marrow. **(A)** Representative flow cytometry plots demonstrating intracellular XBP1s expression in CD11b^+^B220^^-^^CD11c^+^Gr-1^^-^^I-A/I-E^^-^^XBP1s^+^ pre-conventional dendritic cells (pre-cDCs) within fBM. **(B)** Absolute numbers of XBP1s^+^ pre-cDCs in BM of fetuses from OM-85-treated and untreated mothers. Data are presented from individual animals comparing fetuses from OM-85-treated and untreated mothers and displayed as bar graphs showing mean ± SEM of *n* = 4 independent experiments. Statistical significance was determined using Mann-Whitney *U* test.

## Discussion

In this study, we have characterised the response of the fBM myeloid cell compartment to maternal treatment with the microbial-derived innate immune modulator OM-85 during pregnancy. We demonstrate that, consistent with our previous reports in 6-week-old offspring^15^, maternal OM-85 treatment expanded the baseline pool of GMP and MDP in fBM. Furthermore, we show for the first time the maternal OM-85-induced modulation of metabolic pathways within fBM responsible for cellular cholesterol homeostasis, with specific activation of SREBF1, SREBF2, SCAP and LDLR. In this regard, SREBF1 and SREBF2 are responsible for encoding sterol regulatory element binding proteins (SREBP) which play a central role in cellular metabolism by controlling the synthesis of cholesterol and other membrane lipids in the Golgi of mammalian cells, with SREBP2 (encoded by SREBF2) regarded as the “master regulator” of cellular cholesterol biosynthesis^32^. Furthermore, SCAP is central to this process by acting as a protein chaperone, mediating the transport of SREBP from the ER to the Golgi to promote cholesterol biosynthesis^32^. In parallel with this metabolic pathway, LDLR regulates cellular uptake of low-density lipoprotein (LDL), the upregulation of which increases downstream cholesterol accumulation within the cell and is required for the proliferation of hematopoietic stem and progenitor cells^33^. It is now recognised that myelopoiesis within the BM is heavily reliant upon an increased demand for cellular cholesterol and enhanced cholesterol biosynthesis^34-36^, and we therefore postulate that upregulation of this immunometabolic pathway following maternal OM-85 treatment is in-part responsible for the expansion of MP, GMP and MDP observed within fBM. Further reinforcing the importance of enhanced cholesterol biosynthesis, INSIG1, an ER membrane protein that prevents trafficking of SCAP/SREBP complexes to the Golgi and thereby restrains cholesterol biosynthesis^37^, was inhibited in fBM following maternal OM-85 treatment. Together, these findings parallel previous studies demonstrating that enhanced cellular cholesterol biosynthesis and resultant expansion of BM MP is a hallmark of classical β-glucan-mediated central immune training^10^. We further demonstrate that transplacental mechanisms promoting maternal OM-85-induced immune training within fBM involve a dynamic process comprising upregulation of immunometabolic pathways that provide key rate-limiting metabolites required for myelopoiesis and subsequent expansion of MP, GMP and MDP, and concomitant inhibition of negative feedback loops responsible for arresting cholesterol biosynthesis.

Downstream of the fBM progenitor response, maternal OM-85 treatment selectively amplified the overall abundance of fBM cDC, along with enhancing the concomitant functional maturation of these cDC as demonstrated by upregulated I-A/I-E (MHC Class II) expression. This BM population is the source of the precursors which subsequently seed the airway mucosal DC network that progressively develops between birth and weaning^3, 4^. It is noteworthy that the cDC which initially seed this network postnatally are MHC-II^low^ (reflecting their functionally immature status) relative to the high-level expression seen at later ages^3^, and the findings above in fetal cDC from the treated group may collectively explain the accelerated postnatal establishment and the enhanced functional maturation of this network observed in their offspring^15^. This DC network plays an essential “gatekeeper” role in immune surveillance of airway surfaces, and hence in protection against both allergic and infectious diseases in the respiratory tract^38, 39^, and its relative paucity and reduced functionality during infancy may be an important contributor to increased susceptibility to these diseases during this life phase.

Earlier studies from our group also identified unique features of the population dynamics of this lung cDC network which distinguishes it from comparable populations in other tissues, notably the exceptionally rapid baseline turnover rate of individual cells within the network, ~85% of which are replaced every 24-36 hours^40^, with emigration to draining lymph nodes (bearing samples of locally acquired antigens) balanced by recruitment of replacements from BM. Moreover, once development of functional competence is complete^4^, this network develops capability for rapid expansion to up to 5-fold baseline density in the face of acute challenge with airborne pro-inflammatory irritant, allergenic or microbial stimuli^41^, the latter response exhibiting kinetics that rival neutrophils^42, 43^. These unique population dynamics suggests that even at baseline, mechanisms that promote lung cDC survival are likely to play a crucial role in the capacity of the network to perform its immune surveillance functions which require onward migration to downstream lymph nodes and subsequent interaction with T-cells as opposed to antigen presentation *in situ^38^*; moreover during prolonged/severe events exemplified by severe viral infections, the added effects of cDC injury resulting from inflammation-associated ER stress^44^ would place further pressure on survival times.

In this regard, our transcriptomic analyses of fBM from offspring of OM-85-treated mothers also identified upregulated expression of XBP1, ATF6β and ERN1 (encoding IRE1α), key components of the XBP1-ERN1 signalling axis and critical regulators of the UPR pathway^45^ which mitigates the effects of ER stress, and moreover we localised upregulated production of active XBP1s protein to cDC precursors. Taken together with recent findings on the role for IRE1α-XBP1 signalling and the downstream transcription factor XBP1s in DC development and function^22, 23^, these results suggest a central role for the XBP1-ERN1 signalling axis in this OM-85-mediated immune training process.

In further support of this suggestion, other studies demonstrate a significant reduction in CD11c^+^ cells (mirroring that of our pre-cDC phenotype) within XBP1^-/-^ BM cultures, whilst forced overexpression of XBP1s in XBP1^-/-^ DC precursors conversely rescues and subsequently drives expansion of the DC pool *in vitro^22^.* It is also pertinent to note that others have reported that the cDC population in the lung mucosa is differentially reliant upon XBP1 expression for survival at baseline relative to cDC from other tissue sites^44^, which may be a direct reflection of the uniquely high turnover rates of cDC within the airway mucosal microenvironment^40^.

Collectively, the data presented here indicate that OM-85 likely operates as an immune training agent, employing cellular and immunometabolic mechanisms previously reported in independent model systems^10, 11, 46^, with the additional capacity to act transplacentally via the fBM. Furthermore, we go beyond the currently known features of innate immune training to describe involvement of the XBP1-ERN1 signalling axis. Moreover, classical β-glucan- and BCG-mediated immune training has traditionally focused on prototypic innate effector cell populations (monocytes/macrophages/natural killer cells) resulting in enhanced resistance to bystander pathogens via the upregulation of pro-inflammatory responses exemplified by tumour necrosis factor-a (TNF-a), interferon-g (IFN-g), IL-6 and IL-1β production^47-49^ and emergency granulopoiesis-mediated neutrophilic influx^46^. However, studies are now beginning to recognise that immune training can also occur in DC populations, resulting *inter alia* in epigenetic reprograming of pro-inflammatory cytokine responses^50^. We have recently extended these observations to include OM-85-mediated training effects on key immunoregulatory functions in both pregnant mice and their offspring, including effects on both cDC and pDC populations and downstream T-regulatory cells, which are collectively associated with enhanced resistance of both mothers and offspring to the pro-inflammatory effects of bacterial, viral and allergenic stimulation^15, 51^. Of note, similar transplacental immune training-like effects targeting immunoregulatory mechanisms in offspring have been reported in relation to pregnant maternal exposure to extracts from *Acinetobacter lwoffii*^52^ and *Helicobacter pylori*^53^, suggesting that the phenomenon reported here may be generalisable, and may point towards a novel approach to mitigation of disease risk in the age group that is in greatest need of protection.

## Methods

### Animals

Specific pathogen-free BALB/c mice were purchased from the Animal Resource Centre (Murdoch, Western Australia, Australia). All mice were housed under specific pathogen-free conditions at the Telethon Kids Institute Bioresources Centre.

### Time-mated pregnancies

Female BALB/c mice 8-12 weeks of age were time-mated with male BALB/c studs 8-26 weeks of age. Male studs were housed individually with 1-2 females overnight. The detection of a vaginal plug the following morning was designated gestation day (GD) 0.5.

### Maternal OM-85 treatment

OM-85 (OM Pharma) is an endotoxin-low lyophilised extract containing a cocktail of TLR ligands derived from 8 major respiratory tract bacterial pathogens *(Haemophilus influenzae, Streptococcus pneumoniae, Streptococcus pyogenes, Streptococcus viridians, Klebsiella pneumoniae, Klebsiella ozaenae, Staphylococcus aureus* and *Neisseria catarrhalis*)^54, 55^. Based on previously optimised dosing concentrations^51, 56^, time-mated pregnant BALB/c mice selected at random received daily oral feeding of lyophilised OM-85 reconstituted in phosphate-buffered saline (PBS) to a concentration of 400mg/kg body weight for the second half of gestation (GD9.5 – 17.5). Control pregnant mice were left untreated for the duration of the study. All maternal treatment was performed with a single batch of OM-85 (batch 1812162).

### Tissue collection

Pregnant BALB/c mice were sacrificed 24 hours after the final OM-85 dose at GD18.5. Both horns of the uterus were removed and fetuses sacrificed by decapitation. Fetal hind legs (cleaned of excess tissue) were removed. Fetal hind leg samples for flow cytometry were collected into cold PBS + 0.1% bovine serum albumin (BSA) and stored on ice. Fetal hind leg samples for transcriptomic analysis were collected into RNAlater^®^ stabilisation solution (Ambion). Samples collected into RNAlater^®^ were stored overnight at 4°C, then transferred to 1.5ml Eppendorf tubes (Eppendorf) and frozen at −80°C for future transcriptome profiling. Dead fetuses were excluded from the study.

### Single-cell suspension preparation

Fetal hind legs were prepared by mincing with a scalpel followed by enzymatic digestion, as previously detailed^15^. Briefly, minced tissue was resuspended in 10ml GKN + 10% fetal calf serum (FCS; Serana) with collagenase IV (Worthington Biochemical Corp.) and DNase (Sigma-Aldrich) at 37°C under gentle agitation for 60 minutes. Digested cells were filtered through sterile cotton wool columns, centrifuged and resuspended in cold PBS for total cell counts.

### Flow cytometry

Fetal bone marrow single-cell suspensions were used for all immunostaining. Panels of monoclonal antibodies were developed to enable phenotypic characterisation of committed myeloid cells: CD3-FITC (BD Bioscience; clone 17A2), CD11b-BV510 (BD Bioscience; clone M1/70), CD11c-BV711 (BD Bioscience; clone HL3), CD19-APC-H7 (BD Bioscience; clone 1D3), Gr-1-Biotin (BD Bioscience; clone RB6-6B2), CD45R/B220-PerCP-Cy5.5 (BD Bioscience; clone RA3-6B2) NKp46-PE-Cy7 (BioLegend; clone 29A1.4), SIRP-a-APC (BioLegend; clone P84), I-A/I-E-BV421 (BioLegend; clone M5/114.15.2), F4/80-BV785 (BioLegend; clone BM8), Viability-AF700 (BD Bioscience), Streptavidin-BV605 (BD Bioscience); hematopoietic stem and progenitor cells: CD2-Biotin (BD Bioscience; clone RM2-5), CD3-Biotin (BD Bioscience; clone 145-2C11), CD4-Biotin (BD Bioscience; clone GK1.5), CD5-Biotin (BD Bioscience; clone 53-7.3), CD8α-Biotin (BD Bioscience; clone 53-6.7), CD19-Biotin (BD Bioscience; clone 1D3), CD45R/B220-Biotin (BD Bioscience; RA3-6B2), Gr-1-Biotin (BD Bioscience; clone RB6-6B2), Ter119-Biotin (BD Bioscience; clone TER-119), CD16/32-PerCP-Cy5.5 (BD Bioscience; clone 2.4G2), CD34-FITC (BD Bioscience; clone RAM34), IL-7Ra-PE-Cy7 (BD Bioscience; clone SB/199), Flt-3-PE (BD Bioscience; clone A2F10.1), c-Kit-APC-Cy7 (BD Bioscience; clone 2B8), Sca-1-BV510 (BD Bioscience; clone D7), CX3CR1-APC (BioLegend; clone SA011F11), NKG2D-BV711 (BD Bioscience; CX5), Viability-AF700 (BD Bioscience), Streptavidin-BV605 (BD Bioscience) and XBP1s-expressing bone marrow cells: CD3-FITC (BD Bioscience; clone 17A2), CD11b-BV510 (BD Bioscience; clone M1/70), CD11c-AF700 (BD Bioscience; clone HL3), CD19-APC-H7 (BD Bioscience; clone 1D3), I-A/I-E-AF647 (BD Bioscience; clone M5/114.15.2), CD45R/B220-PE-CF594 (BD Bioscience; RA3-6B2), Gr-1-Biotin (BD Bioscience; clone RB6-6B2), NKp46-PE-Cy7 (BioLegend; clone 29A1.4), F4/80-BV785 (BioLegend; clone BM8), XBP1s-BV421 (BD Bioscience; clone Q3-695), Streptavidin-BV605 (BD Bioscience). Intracellular staining for XBP1s was performed using an intracellular staining buffer kit (eBioscience). Data acquisition was performed on a 4-laser LSR Fortessa (BD Bioscience). All samples were kept as individuals and not pooled. Immune cell phenotypic characterisation was performed using FlowJo software (version 10.1, Tree Star). Fluorescence minus one staining controls were used for all panels.

### Flow cytometric statistical analyses

Statistical analysis and graphing was performed using GraphPad Prism (GraphPad software; version 7.0a). Statistical significance of p<0.05 was considered significant. Unpaired, two-tailed Student’s *t*-test or Mann Whitney *U* test were used based on distribution of the data as determined by D’Agostino-Pearson omnibus normality test. Part of the cDC flow cytometry data presented in Figure 1 and MDP data presented in Figure 2 has been published in a forerunner manuscript^15^.

### Transcriptome profiling (RNA-Seq)

*Tissue preparation, RNA extraction and transcriptome profiling.* Fetal bone marrow samples were homogenised using a rotor-star homogeniser (Qiagen) and total RNA extracted via TRIzol (Ambion), followed by clean-up using RNeasy MinElute (Qiagen). RNA integrity was determined using an Agilent 2100 Bioanalyzer (Agilent Technologies; RIN: 10 ± 0 (mean ± SD)). 1μg total RNA (*n* = 32) was shipped on dry ice to the Australia Genome Research Facility (AGRF) for library preparation (TruSeq Stranded mRNA Library Prep Kit, Illumina) and sequencing (Illumina HiSeq2500, 50-bp singe-end reads, v4 chemistry). The raw sequencing data are available from GEO (accession; GSE140143).

## RNA-Seq data analysis

*Pre-processing and exploratory data analysis:* RNA-Seq data was analysed in the R environment for statistical computing. Sequencing data quality control (QC) was performed with the Bioconductor package Rqc^57^. Sequencing reads were aligned to the reference murine genome (mm10) using Subread and summarised at the gene level using featureCounts^58^. Genes with <500 total counts across the data were removed from the analysis. Sample QC was performed by analysing the distribution of the raw read counts to check for sample outliers using boxplots, relative log-transformed expression (RLE) plots and principal component analysis (PCA), before and after global-scale median normalisation. *Differential expression analysis:* Differentially expressed genes (DEG) were identified employing the *DESeq2* package^59^. *DESeq2* utilises a negative binomial distribution model, with a False Discovery Rate adjusted *P*-value for multiple comparisons. Genes were deemed significant with an adjusted *P*-value < 0.1. *Pathways analysis:* Pathways enrichment analysis was performed using the InnateDB database^16^ with Benjamini & Hochberg adjusted *P*-value ≤ 0.05 deemed significant. *Upstream regulator analysis:* Ingenuity Systems Upstream Regulator Analysis^60^ was employed to identify putative molecular drives of the DEG patterns. Significance was determined by activation Z-score ≥ 2 and *P*-value of overlap ≤ 0.05.

## Study approval

All animal experiments were formally approved by the Telethon Kids Institute Animal Ethics Committee, operating under the guidelines developed by the National Health and Medical Research Council of Australia for the care and use of animals in scientific research.

## Acknowledgements

The authors wish to acknowledge the animal technicians at the Telethon Kids Institute Bioresources Centre. This study was funded by the National Health and Medical Research Council of Australia. OM Pharma provided the OM-85 at no cost.

## Author contributions

KTM, PGH and DHS designed the study. KTM, MB, NMS and JFLJ performed the experiments. KTM, ACJ, MB and DHS analysed the data. PAS and AB contributed to the project design and discussions on data interpretation. KTM, PAS, PGH and DHS wrote the manuscript. All authors reviewed the final version of the manuscript.

