## Supplementary Data for "Transplacental innate immune training via maternal microbial exposure: the XBP1-ERN1 axis in programming dendritic cell precursors"

**Supplementary Table 1. Differentially expressed genes comparing bone marrow of fetuses from OM-85 treated versus untreated mothers.**

| GeneID | Symbol | baseMean | log2FC | lfcSE | stat | p-value | adjusted p-value |
|---|---|---|---|---|---|---|---|
| 231070 | Insig1 | 1227.87 | 0.31 | 0.08 | 3.76 | 1.73E-04 | 5.99E-02 |
| 319554 | Idi1 | 297.74 | 0.29 | 0.07 | 3.99 | 6.63E-05 | 4.00E-02 |
| 78894 | Aacs | 1184.93 | 0.28 | 0.07 | 4.23 | 2.32E-05 | 2.66E-02 |
| 64136 | Sdf2l1 | 470.14 | 0.28 | 0.08 | 3.58 | 3.38E-04 | 7.01E-02 |
| 240025 | Dact2 | 1114.16 | 0.28 | 0.08 | 3.35 | 7.97E-04 | 9.05E-02 |
| 16835 | Ldlr | 2953.52 | 0.27 | 0.07 | 3.67 | 2.47E-04 | 6.78E-02 |
| 71911 | Bdh1 | 914.76 | 0.27 | 0.08 | 3.34 | 8.45E-04 | 9.05E-02 |
| 19273 | Ptpru | 284.90 | 0.25 | 0.08 | 3.25 | 1.14E-03 | 9.46E-02 |
| 331026 | Gmppb | 548.39 | 0.24 | 0.06 | 3.95 | 7.92E-05 | 4.11E-02 |
| 56325 | Abcb9 | 518.14 | 0.24 | 0.07 | 3.56 | 3.77E-04 | 7.01E-02 |
| 76737 | Creld2 | 834.95 | 0.24 | 0.05 | 4.72 | 2.40E-06 | 1.37E-02 |
| 74246 | Gale | 1123.69 | 0.23 | 0.06 | 4.20 | 2.67E-05 | 2.66E-02 |
| 71609 | Tradd | 312.97 | 0.23 | 0.06 | 4.16 | 3.24E-05 | 2.66E-02 |
| 192156 | Mvd | 666.21 | 0.23 | 0.06 | 3.71 | 2.10E-04 | 6.48E-02 |
| 59031 | Chst12 | 813.08 | 0.22 | 0.07 | 3.38 | 7.32E-04 | 9.00E-02 |
| 74840 | Manf | 2133.29 | 0.22 | 0.05 | 4.35 | 1.36E-05 | 2.32E-02 |
| 17428 | Mnt | 1032.31 | 0.21 | 0.06 | 3.49 | 4.80E-04 | 7.50E-02 |
| 233912 | Armc5 | 603.81 | 0.20 | 0.06 | 3.61 | 3.03E-04 | 7.01E-02 |
| 13360 | Dhcr7 | 1513.17 | 0.20 | 0.06 | 3.33 | 8.54E-04 | 9.05E-02 |
| 15015 | H2-Q4 | 790.97 | 0.19 | 0.06 | 3.27 | 1.08E-03 | 9.24E-02 |
| 20787 | Srebf1 | 4157.10 | 0.19 | 0.05 | 3.49 | 4.87E-04 | 7.50E-02 |
| 68671 | Pcyt2 | 1658.43 | 0.19 | 0.04 | 4.81 | 1.48E-06 | 1.37E-02 |
| 72296 | Rusc1 | 541.78 | 0.19 | 0.05 | 3.84 | 1.21E-04 | 5.11E-02 |
| 56470 | Rgs19 | 955.91 | 0.18 | 0.04 | 4.38 | 1.21E-05 | 2.32E-02 |
| 230145 | Galnt12 | 574.23 | 0.18 | 0.05 | 3.92 | 8.80E-05 | 4.37E-02 |
| 56473 | Fads2 | 1749.89 | 0.18 | 0.05 | 3.79 | 1.51E-04 | 5.76E-02 |
| 235293 | Sc5d | 1018.49 | 0.18 | 0.05 | 3.76 | 1.67E-04 | 5.96E-02 |
| 320534 | Tmem104 | 400.07 | 0.18 | 0.05 | 3.40 | 6.72E-04 | 8.62E-02 |
| 13358 | Slc25a1 | 769.68 | 0.18 | 0.05 | 3.23 | 1.23E-03 | 9.83E-02 |
| 12317 | Calr | 9484.29 | 0.18 | 0.05 | 3.65 | 2.62E-04 | 6.78E-02 |
| 67065 | Polr3d | 801.45 | 0.17 | 0.05 | 3.67 | 2.40E-04 | 6.78E-02 |
| 225849 | Ppp2r5b | 654.98 | 0.17 | 0.05 | 3.56 | 3.73E-04 | 7.01E-02 |
| 12915 | Atf6b | 1644.33 | 0.17 | 0.05 | 3.35 | 8.16E-04 | 9.05E-02 |
| 228368 | Slc35c1 | 1022.13 | 0.17 | 0.04 | 4.20 | 2.70E-05 | 2.66E-02 |
| 18799 | Plcd1 | 1381.75 | 0.17 | 0.05 | 3.36 | 7.73E-04 | 9.00E-02 |
| 103724 | Tbc1d10a | 827.76 | 0.17 | 0.05 | 3.43 | 5.99E-04 | 8.00E-02 |
| 109815 | Vimp | 368.39 | 0.16 | 0.05 | 3.25 | 1.15E-03 | 9.46E-02 |
| 57377 | Mogs | 2743.90 | 0.16 | 0.04 | 3.65 | 2.62E-04 | 6.78E-02 |
| 64144 | Mllt1 | 2427.05 | 0.16 | 0.05 | 3.30 | 9.83E-04 | 9.16E-02 |

| GeneID | Symbol | baseMean | log2FC | lfcSE | stat | p-value | adjusted p-value |
|---|---|---|---|---|---|---|---|
| 26394 | Lypla2 | 1464.47 | 0.16 | 0.05 | 3.46 | 5.36E-04 | 7.80E-02 |
| 72056 | 1810055G02Rik | 1035.38 | 0.16 | 0.04 | 3.72 | 2.02E-04 | 6.46E-02 |
| 72029 | Cnpy3 | 1174.74 | 0.16 | 0.04 | 4.25 | 2.16E-05 | 2.66E-02 |
| 103425 | Ncln | 2243.69 | 0.16 | 0.05 | 3.31 | 9.35E-04 | 9.14E-02 |
| 269881 | Map3k10 | 498.33 | 0.16 | 0.05 | 3.38 | 7.25E-04 | 9.00E-02 |
| 74126 | Syvn1 | 1907.98 | 0.16 | 0.05 | 3.22 | 1.29E-03 | 9.99E-02 |
| 23991 | Cib1 | 731.86 | 0.15 | 0.04 | 3.65 | 2.67E-04 | 6.78E-02 |
| 73062 | Ppp1r16a | 739.65 | 0.15 | 0.04 | 3.87 | 1.09E-04 | 4.80E-02 |
| 68017 | Mrm2 | 213.39 | 0.15 | 0.05 | 3.28 | 1.04E-03 | 9.18E-02 |
| 93685 | Entpd7 | 521.74 | 0.15 | 0.04 | 3.33 | 8.72E-04 | 9.05E-02 |
| 107242 | AI837181 | 1237.54 | 0.15 | 0.04 | 3.54 | 3.93E-04 | 7.01E-02 |
| 23917 | Impdh1 | 2864.39 | 0.15 | 0.04 | 3.36 | 7.69E-04 | 9.00E-02 |
| 12304 | Pdia4 | 4125.69 | 0.14 | 0.04 | 3.28 | 1.03E-03 | 9.18E-02 |
| 270066 | Slc35e1 | 2671.84 | 0.14 | 0.03 | 4.42 | 9.91E-06 | 2.32E-02 |
| 277010 | Marveld1 | 2058.69 | 0.14 | 0.04 | 3.21 | 1.33E-03 | 9.99E-02 |
| 70314 | Rabep2 | 533.66 | 0.14 | 0.04 | 3.52 | 4.32E-04 | 7.01E-02 |
| 67789 | Dalrd3 | 865.66 | 0.14 | 0.04 | 3.21 | 1.31E-03 | 9.99E-02 |
| 234378 | Klhl26 | 658.37 | 0.14 | 0.04 | 3.28 | 1.03E-03 | 9.18E-02 |
| 223690 | Ankrd54 | 788.15 | 0.14 | 0.04 | 3.55 | 3.81E-04 | 7.01E-02 |
| 231807 | BC037034 | 1012.51 | 0.14 | 0.04 | 3.63 | 2.89E-04 | 7.01E-02 |
| 664994 | Isoc2a | 457.74 | 0.14 | 0.04 | 3.65 | 2.60E-04 | 6.78E-02 |
| 71853 | Pdia6 | 7221.20 | 0.14 | 0.04 | 3.59 | 3.26E-04 | 7.01E-02 |
| 18081 | Ninj1 | 439.39 | 0.14 | 0.04 | 3.21 | 1.32E-03 | 9.99E-02 |
| 208638 | Slc25a38 | 505.11 | 0.14 | 0.04 | 3.29 | 9.98E-04 | 9.16E-02 |
| 22433 | Xbp1 | 3609.25 | 0.14 | 0.04 | 3.58 | 3.47E-04 | 7.01E-02 |
| 26433 | Plod3 | 2737.30 | 0.14 | 0.04 | 3.23 | 1.22E-03 | 9.83E-02 |
| 56214 | Scamp4 | 1189.97 | 0.14 | 0.04 | 3.60 | 3.22E-04 | 7.01E-02 |
| 66058 | Tmem176a | 1019.93 | 0.14 | 0.04 | 3.29 | 9.94E-04 | 9.16E-02 |
| 20818 | Srprb | 1450.32 | 0.14 | 0.04 | 3.54 | 3.98E-04 | 7.01E-02 |
| 503610 | Zdhhc18 | 1356.49 | 0.14 | 0.04 | 3.57 | 3.61E-04 | 7.01E-02 |
| 70231 | Gorasp2 | 3387.38 | 0.13 | 0.04 | 3.46 | 5.40E-04 | 7.80E-02 |
| 106200 | Txndc11 | 934.54 | 0.13 | 0.03 | 3.99 | 6.65E-05 | 4.00E-02 |
| 74451 | Pgs1 | 876.13 | 0.13 | 0.03 | 3.89 | 1.00E-04 | 4.73E-02 |
| 68427 | Slc39a13 | 1519.83 | 0.13 | 0.04 | 3.37 | 7.47E-04 | 9.00E-02 |
| 217664 | Mgat2 | 1853.11 | 0.13 | 0.04 | 3.45 | 5.58E-04 | 7.85E-02 |
| 22232 | Slc35a2 | 1120.66 | 0.13 | 0.04 | 3.52 | 4.36E-04 | 7.01E-02 |
| 22687 | Zpr1 | 1098.39 | 0.13 | 0.03 | 4.03 | 5.50E-05 | 3.69E-02 |
| 100090 | Zbtb48 | 459.73 | 0.13 | 0.04 | 3.26 | 1.11E-03 | 9.42E-02 |
| 76267 | Fads1 | 3584.00 | 0.13 | 0.03 | 4.34 | 1.42E-05 | 2.32E-02 |
| 71116 | Stx18 | 603.81 | 0.13 | 0.03 | 4.14 | 3.50E-05 | 2.66E-02 |
| 218271 | B4galt7 | 571.18 | 0.13 | 0.04 | 3.29 | 1.00E-03 | 9.16E-02 |

| GeneID | Symbol | baseMean | log2FC | lfcSE | stat | p-value | adjusted p-value |
|---|---|---|---|---|---|---|---|
| 53421 | Sec61a1 | 6887.06 | 0.12 | 0.03 | 3.77 | 1.66E-04 | 5.96E-02 |
| 68385 | Tlcd1 | 319.70 | 0.12 | 0.04 | 3.31 | 9.38E-04 | 9.14E-02 |
| 20405 | Sh3gl1 | 2885.28 | 0.12 | 0.04 | 3.43 | 6.02E-04 | 8.00E-02 |
| 54399 | Bet1l | 782.18 | 0.12 | 0.03 | 3.53 | 4.11E-04 | 7.01E-02 |
| 76025 | Cant1 | 1650.55 | 0.12 | 0.03 | 3.96 | 7.36E-05 | 4.03E-02 |
| 20514 | Slc1a5 | 1765.32 | 0.12 | 0.03 | 3.43 | 6.08E-04 | 8.00E-02 |
| 11993 | Aup1 | 1989.09 | 0.12 | 0.03 | 3.47 | 5.29E-04 | 7.80E-02 |
| 28106 | Mydgf | 1263.52 | 0.12 | 0.03 | 3.37 | 7.63E-04 | 9.00E-02 |
| 20498 | Slc12a4 | 2174.73 | 0.12 | 0.04 | 3.31 | 9.44E-04 | 9.14E-02 |
| 20832 | Ssr4 | 1760.63 | 0.12 | 0.03 | 3.53 | 4.12E-04 | 7.01E-02 |
| 19246 | Ptpn1 | 1994.99 | 0.12 | 0.03 | 3.45 | 5.64E-04 | 7.85E-02 |
| 66357 | Ostc | 2242.03 | 0.12 | 0.03 | 3.57 | 3.62E-04 | 7.01E-02 |
| 72727 | B3gat3 | 1046.49 | 0.11 | 0.04 | 3.22 | 1.26E-03 | 9.94E-02 |
| 56530 | Cnpy2 | 1775.08 | 0.11 | 0.03 | 3.48 | 4.93E-04 | 7.50E-02 |
| 66059 | Krtcap2 | 1263.12 | 0.11 | 0.03 | 3.37 | 7.43E-04 | 9.00E-02 |
| 74504 | Fam53a | 728.64 | 0.11 | 0.03 | 3.26 | 1.11E-03 | 9.42E-02 |
| 73836 | Slc35b2 | 1774.75 | 0.11 | 0.03 | 3.53 | 4.23E-04 | 7.01E-02 |
| 52858 | Cdipt | 1544.68 | 0.11 | 0.03 | 4.14 | 3.43E-05 | 2.66E-02 |
| 17308 | Mgat1 | 2619.94 | 0.11 | 0.03 | 3.62 | 2.96E-04 | 7.01E-02 |
| 66156 | Anapc11 | 977.77 | 0.11 | 0.03 | 3.31 | 9.30E-04 | 9.14E-02 |
| 56457 | Clptm1 | 2346.11 | 0.11 | 0.03 | 3.44 | 5.89E-04 | 8.00E-02 |
| 68047 | Mpnd | 1339.91 | 0.11 | 0.03 | 3.42 | 6.21E-04 | 8.06E-02 |
| 71667 | Tmem248 | 1805.98 | 0.10 | 0.03 | 3.45 | 5.58E-04 | 7.85E-02 |
| 13852 | Stx2 | 934.84 | 0.10 | 0.03 | 3.53 | 4.16E-04 | 7.01E-02 |
| 20932 | Surf4 | 8479.03 | 0.10 | 0.03 | 3.71 | 2.04E-04 | 6.46E-02 |
| 68090 | Yif1a | 900.31 | 0.10 | 0.03 | 3.55 | 3.79E-04 | 7.01E-02 |
| 14792 | Lpcat3 | 1414.89 | 0.10 | 0.03 | 4.07 | 4.72E-05 | 3.37E-02 |
| 76479 | Smndc1 | 1143.61 | 0.10 | 0.03 | 3.33 | 8.60E-04 | 9.05E-02 |
| 84095 | Pi4k2a | 1255.74 | 0.10 | 0.03 | 3.27 | 1.07E-03 | 9.24E-02 |
| 67511 | Tmed9 | 2708.52 | 0.10 | 0.03 | 3.83 | 1.28E-04 | 5.23E-02 |
| 11867 | Arpc1b | 2894.10 | 0.10 | 0.03 | 3.48 | 5.07E-04 | 7.62E-02 |
| 236732 | Rbm10 | 1178.82 | 0.10 | 0.03 | 3.24 | 1.18E-03 | 9.58E-02 |
| 20333 | Sec22b | 2051.88 | 0.10 | 0.03 | 3.43 | 6.09E-04 | 8.00E-02 |
| 12313 | Calm1 | 10795.59 | 0.09 | 0.03 | 3.33 | 8.68E-04 | 9.05E-02 |
| 72055 | Slc38a10 | 4184.42 | 0.09 | 0.03 | 3.37 | 7.60E-04 | 9.00E-02 |
| 64143 | Ralb | 880.29 | 0.09 | 0.03 | 3.23 | 1.25E-03 | 9.90E-02 |
| 68944 | Tmco1 | 877.50 | 0.09 | 0.03 | 3.52 | 4.32E-04 | 7.01E-02 |
| 11848 | Rhoa | 4109.43 | 0.08 | 0.02 | 3.32 | 8.92E-04 | 9.14E-02 |
| 20529 | Slc31a1 | 1843.88 | 0.07 | 0.02 | 3.33 | 8.64E-04 | 9.05E-02 |
| 75717 | Cul5 | 1551.18 | -0.08 | 0.02 | -3.49 | 4.77E-04 | 7.50E-02 |
| 19317 | Qk | 6579.70 | -0.09 | 0.03 | -3.33 | 8.60E-04 | 9.05E-02 |

| GeneID | Symbol | baseMean | log2FC | lfcSE | stat | p-value | adjusted p-value |
|---|---|---|---|---|---|---|---|
| 380916 | Lrch1 | 1181.85 | -0.10 | 0.03 | -3.70 | 2.18E-04 | 6.54E-02 |
| 59125 | Nek7 | 1619.69 | -0.11 | 0.03 | -3.73 | 1.91E-04 | 6.40E-02 |
| 81003 | Trim23 | 702.07 | -0.12 | 0.03 | -3.60 | 3.17E-04 | 7.01E-02 |
| 381306 | BC055324 | 684.71 | -0.13 | 0.04 | -3.22 | 1.28E-03 | 9.99E-02 |
| 14594 | Ggta1 | 2022.88 | -0.13 | 0.04 | -3.34 | 8.33E-04 | 9.05E-02 |
| 103573 | Xpo1 | 8600.19 | -0.13 | 0.04 | -3.25 | 1.14E-03 | 9.46E-02 |
| 20843 | Stag2 | 6488.16 | -0.14 | 0.04 | -3.32 | 9.03E-04 | 9.14E-02 |
| 433931 | Pigg | 382.49 | -0.14 | 0.04 | -3.28 | 1.05E-03 | 9.18E-02 |
| 245631 | Mum1l1 | 1048.22 | -0.15 | 0.04 | -3.34 | 8.26E-04 | 9.05E-02 |
| 26399 | Map2k6 | 453.63 | -0.15 | 0.04 | -3.34 | 8.37E-04 | 9.05E-02 |
| 211329 | Ncoa7 | 568.38 | -0.15 | 0.04 | -3.69 | 2.24E-04 | 6.56E-02 |
| 329260 | Dennd1b | 874.05 | -0.15 | 0.05 | -3.32 | 9.14E-04 | 9.14E-02 |
| 76967 | 2700049A03Rik | 775.57 | -0.15 | 0.05 | -3.25 | 1.15E-03 | 9.46E-02 |
| 18583 | Pde7a | 1257.44 | -0.15 | 0.04 | -3.81 | 1.37E-04 | 5.38E-02 |
| 78796 | Zcchc4 | 234.07 | -0.16 | 0.05 | -3.28 | 1.04E-03 | 9.18E-02 |
| 17957 | Napb | 219.35 | -0.16 | 0.05 | -3.21 | 1.30E-03 | 9.99E-02 |
| 230259 | E130308A19Rik | 261.16 | -0.16 | 0.05 | -3.39 | 7.11E-04 | 9.00E-02 |
| 229473 | D930015E06Rik | 5508.16 | -0.17 | 0.04 | -4.15 | 3.28E-05 | 2.66E-02 |
| 54598 | Calcrl | 1124.86 | -0.18 | 0.06 | -3.29 | 9.87E-04 | 9.16E-02 |
| 14007 | Celf2 | 5372.06 | -0.19 | 0.06 | -3.29 | 1.00E-03 | 9.16E-02 |
| 54610 | Tbc1d8 | 732.43 | -0.20 | 0.05 | -3.56 | 3.76E-04 | 7.01E-02 |
| 78286 | Nav2 | 3091.40 | -0.21 | 0.06 | -3.54 | 3.98E-04 | 7.01E-02 |
| 12894 | Cpt1a | 2270.83 | -0.23 | 0.06 | -3.96 | 7.42E-05 | 4.03E-02 |
| 14051 | Eya4 | 664.37 | -0.24 | 0.07 | -3.27 | 1.08E-03 | 9.24E-02 |
| 241589 | D430041D05Rik | 864.80 | -0.26 | 0.08 | -3.21 | 1.32E-03 | 9.99E-02 |
| 77963 | Hook1 | 626.22 | -0.27 | 0.08 | -3.24 | 1.18E-03 | 9.58E-02 |
| 72685 | Dnajc6 | 961.70 | -0.28 | 0.08 | -3.31 | 9.41E-04 | 9.14E-02 |
| 17472 | Gbp4 | 1119.76 | -0.32 | 0.10 | -3.30 | 9.53E-04 | 9.14E-02 |
| 12823 | Col19a1 | 243.16 | -0.33 | 0.09 | -3.88 | 1.04E-04 | 4.73E-02 |
| 57875 | Angptl4 | 338.30 | -0.35 | 0.10 | -3.59 | 3.34E-04 | 7.01E-02 |
| 11535 | Adm | 179.61 | -0.51 | 0.11 | -4.53 | 5.90E-06 | 2.25E-02 |

**Supplementary Table 2. Top 15 InnateDB pathways of differentially expressed upregulated genes comparing bone marrow of fetuses from OM-85 treated versus untreated mothers.**

| **Pathway Name (No. of genes)** | **Gene Symbols** | **Adjusted *P*-value** |
|---|---|---|
| **Protein processing in endoplasmic reticulum (n=10)** | *Atf6b, Calr, Mogs, Pdia4, Pdia6, Sec61a1, Ssr4, Syvn1, Vimp, Xbp1* | 1.67E-05 |
| **Transport of nucleotide sugars (n=3)** | *Slc35a2, Slc35b2, Slc35c1* | 3.52E-04 |
| **Cholesterol biosynthesis (n=4)** | *Dhcr7, Idi1, Mvd, Sc5d* | 4.27E-04 |
| **Regulation of cholesterol biosynthesis by SREBP (SREBF) (n=5)** | *Dhcr7, Idi1, Insig1, Mvd, Sc5d* | 4.55E-04 |
| **Metabolism of lipids and lipoproteins (n=14)** | *Bdh1, Cdipt, Dhcr7, Fads1, Fads2, Idi1, Insig1, Ldlr, Lpcat3, Mvd, Pcyt2, Pgs1, Pi4k2a, Sc5d* | 4.59E-04 |
| **Unfolded Protein Response (UPR) (n=5)** | *Calr, D17Wsu104e, Syvn1, Xbp1, Yif1a* | 8.65E-04 |
| **SNARE interactions in vesicular transport (n=4)** | *Bet1l, Sec22b, Stx18, Stx2* | 9.36E-04 |
| **Activation of gene expression by SREBF (SREBP) (n=4)** | *Dhcr7, Idi1, Mvd, Sc5d* | 9.48E-04 |
| **Asparagine N-linked glycosylation (n=6)** | *Calr, Gmppb, Mgat1, Mgat2, Mogs, Mvd* | 9.57E-04 |
| **Metabolism of proteins (n=13)** | *Calr, D17Wsu104e, Galnt12, Gmppb, Mgat1, Mgat2, Mogs, Mvd, Sec61a1, Ssr4, Syvn1, Xbp1, Yif1a* | 9.86E-04 |
| **XBP1(S) activates chaperone genes (n=4)** | *D17Wsu104e, Syvn1, Xbp1, Yif1a* | 1.61E-03 |
| **IRE1alpha activates chaperones (n=4)** | *D17Wsu104e, Syvn1, Xbp1, Yif1a* | 1.62E-03 |
| **Metabolism (n=21)** | *B3gat3, B4galt7, Bdh1, Cdipt, Chst12, Dhcr7, Fads1, Fads2, Gale, Idi1, Impdh1, Insig1, Ldlr, Lpcat3, Mvd, Pcyt2, Pgs1, Pi4k2a, Plcd1, Sc5d, Slc35b2* | 2.31E-03 |
| **Glycosaminoglycan biosynthesis (n=3)** | *B3gat3, B4galt7, Chst12* | 2.41E-03 |
| **Post-translational protein modification (n=7)** | *Calr, Galnt12, Gmppb, Mgat1, Mgat2, Mogs, Mvd* | 2.50E-03 |

**Supplementary Table 3. Top 20 upstream regulators of differentially expressed genes comparing bone marrow of fetuses from OM-85 treated versus untreated mothers.**

| Upstream Regulator | Molecule Type | Predicted Activation State | Target Genes | Activation z-score | *P*-value |
|---|---|---|---|---|---|
| **XBP1** | Transcription regulator | **Activated** | *ATF6B, BET1L, CALR, GORASP2, MGAT2, MOGS, PDIA4, PDIA6, SDF2L1, SEC22B, SEC61A1, SREBF1, SRPRB, SSR4, STX18, SURF4, SYVN1, TXNDC11, XBP1, YIF1A* | 4.427 | 3.81E-17 |
| **SCAP** | Other | **Activated** | *AACS, DHCR7, FADS2, IDI1, INSIG1, LDLR, MVD, SC5D, SREBF1* | 2.949 | 1.52E-10 |
| **SREBF2** | Transcription regulator | **Activated** | *AACS, DHCR7, FADS2, IDI1, INSIG1, LDLR, MVD, SC5D, SREBF1* | 2.745 | 2.95E-09 |
| **INSIG1** | Other | **Inhibited** | *AACS, DHCR7, FADS1, FADS2, IDI1, LDLR, LPCAT3, PCYT2, SREBF1* | -2.931 | 5.81E-08 |
| **POR** | Enzyme | **Inhibited** | *BDH1, CPT1A, DHCR7, FADS2, HLA-A, IDI1, INSIG1, LDLR, MVD, SC5D, TMEM176A* | -2.8 | 8.67E-08 |
| **SIRT2** | Transcription regulator | **Activated** | *AACS, DHCR7, IDI1, MVD, SC5D* | 2.236 | 1.52E-07 |
| **SREBF1** | Transcription regulator | **Activated** | *AACS, DHCR7, FADS1, FADS2, IDI1, INSIG1, LDLR, MVD, SC5D, SREBF1* | 3.056 | 1.95E-06 |
| **ERN1** | Kinase | **Activated** | *HOOK1, MVD, SDF2L1, SEC22B, SEC61A1, SURF4, SYVN1, XBP1* | 2.156 | 3.32E-06 |
| **CD38** | Enzyme | **Activated** | *B4GALT7, CHST12, CRELD2, MANF, PDIA6, SDF2L1, XBP1* | 2.588 | 1.48E-04 |
| **NFE2L2** | Transcription regulator | **Activated** | *Calm1, DHCR7, IMPDH1, MOGS, PDIA4, PDIA6, PTPN1, SEC61A1, Slc35a2, SREBF1, XBP1* | 2.111 | 1.68E-04 |
| **miR-874-5p** | Mature microrna | **Inhibited** | *ANGPTL4, FAM53A, GALE, KLHL26, LPCAT3, MARVELD1, MNT, NCLN, SCAMP4, SLC35C1, SLC39A13, TMEM104* | -2.887 | 1.88E-04 |
| **miR-4731-5p** | Mature microrna | **Inhibited** | *ARMC5, BET1L, CALR, DACT2, LYPLA2, MNT, NAV2, NINJ1, QKI, SCAMP4, SH3GL1, SLC35E1, SLC39A13, SYVN1, ZDHHC18* | -2.84 | 3.83E-04 |
| **SLC13A1** | Transporter | **Inhibited** | *GALE, INSIG1, MVD, SDF2L1, XBP1* | -2.236 | 5.65E-04 |
| **IL5** | Cytokine | **Activated** | *CHST12, CRELD2, IDI1, MANF, PDIA6, SDF2L1, SLC1A5, XBP1* | 2.758 | 1.07E-03 |
| **miR-4503** | Mature microrna | **Inhibited** | *C11orf24, IDI1, SC5D, SYVN1* | -2 | 1.62E-03 |
| **miR-6887-3p** | Mature microrna | **Inhibited** | *ARPC1B, BDH1, CNPY2, EYA4, FADS2, IMPDH1, ISOC2, MNT, MOGS, PDE7A, SURF4, TMED9* | -2.309 | 1.72E-03 |
| **SIRT1** | Transcription regulator | **Inhibited** | *CPT1A, LDLR, MGAT1, MGAT2, MLLT1, SEC61A1, SREBF1* | -2.395 | 1.96E-03 |
| **miR-6132** | Mature microrna | **Inhibited** | *ARMC5, IMPDH1, MAP3K10, MNT, NINJ1, SH3GL1, SLC25A38, SLC35B2, SLC39A13, TMCO1, TMED9, TMEM176A* | -3.464 | 2.86E-03 |
| **miR-149-3p** | Mature microrna | **Inhibited** | *ANKRD54, BET1L, C7orf43, CALR, CNPY3, FADS2, KLHL26, MARVELD1, MNT, NAV2, RABEP2, RALB, RUSC1, SCAMP4, SH3GL1, SLC35B2, STX2, TMEM104* | -3.771 | 2.99E-03 |
| **miR-6795-5p** | Mature microrna | **Inhibited** | *BET1L, C11orf68, FADS2, ISOC2, KLHL26, LYPLA2, MNT, MOGS, PI4K2A, POLR3D, RGS19, SCAMP4, SYVN1* | -3.606 | 3.03E-03 |

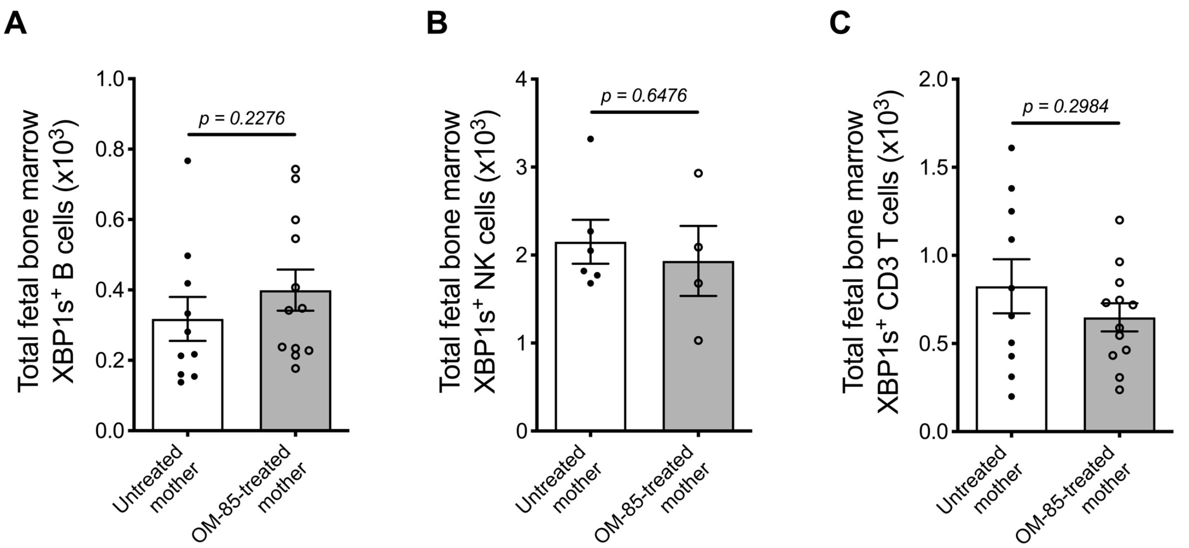


**Supplementary Figure 1. XBP1s is expressed in multiple cell types within fetal bone marrow.** Absolute numbers of **(A)** $CD19^{+}B220^{+}XBP1s^{+}$ B cells, **(B)** $NKp46^{+}CD11b^{+}B220^{+}CD11c^{lo}XBP1s^{+}$ NK cells and **(C)** $CD3^{+}XBP1s^{+}$ T cells in fetal bone marrow. Data are presented from individual animals comparing fetuses from OM-85-treated and untreated mothers and displayed as bar graphs showing mean ± SEM of $n$ = 4 independent experiments. Statistical significance was determined using Mann-Whitney $U$ test (A, B) or Student's $t$ test (C) based on distribution of the data as determined by D'Agostino-Pearson omnibus normality test.
